# Systematic comparison of culture media uncovers phenotypic shift of human microglia defined by reduced reliance to CSF1R signaling

**DOI:** 10.1101/2022.07.14.500101

**Authors:** Marie-France Dorion, Moein Yaqubi, Hunter J. Murdoch, Jeffery A. Hall, Roy Dudley, Jack P. Antel, Thomas M. Durcan, Luke M. Healy

**Author notes:** **Data availability statement:** Sequencing data will be deposited to Gene Expression Omnibus upon acceptance of this manuscript for publication.

## Abstract

Efforts to understand microglia function in health and diseases have been hindered by the lack of culture models that recapitulate *in situ* cellular properties. In recent years, the use of serum-free media with brain-derived growth factors (CSF1R ligands and TGF-β1/2) have been favored for the maintenance of rodent microglia as they promote morphological features observed *in situ*. Here we study the functional and transcriptomic impacts of such media on human microglia. Media formulation had little impact on microglia transcriptome assessed by RNA sequencing which was sufficient to significantly alter microglia capacity to phagocytose myelin debris and to elicit an inflammatory response to lipopolysaccharide. When compared to immediately *ex vivo* microglia from the same donors, the addition of fetal bovine serum to culture media, but not growth factors, was found to aid in the maintenance of key signature genes including those involved in phagocytic processes. A phenotypic shift characterized by *CSF1R* downregulation in culture correlated with a lack of reliance on CSF1R signaling for survival. Consequently, no improvement in cell survival was observed following culture supplementation with CSF1R ligands. Our study provides better understanding of human microglia in culture, with observations that diverge from those previously made in rodent microglia.

**Graphical abstract:** 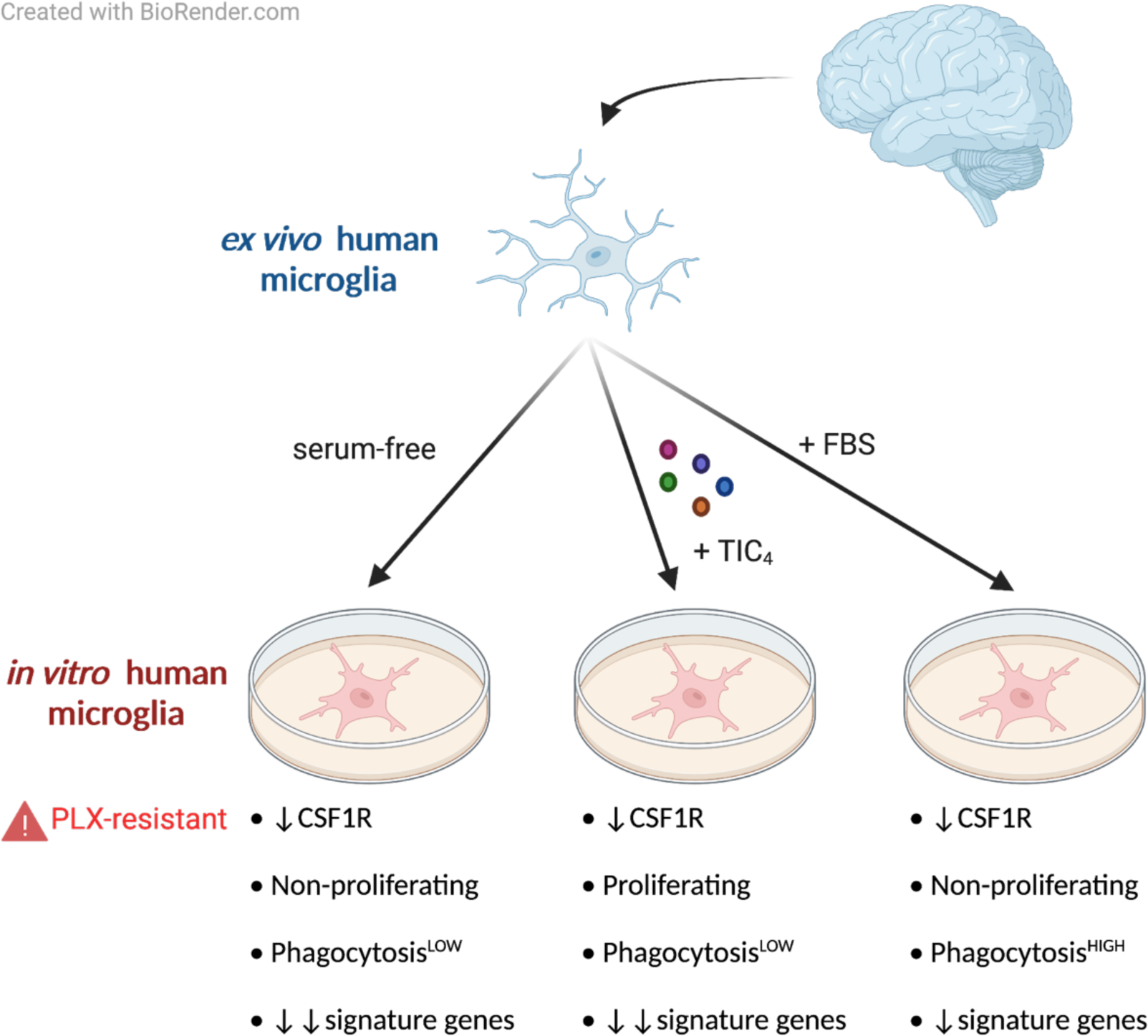

## 1. Introduction

Microglia are highly plastic cells that can undergo changes in their secretory, migratory and phagocytic activities, often accompanied by morphological alterations (Butovsky & Weiner, 2018). In the homeostatic brain, they display a ramified morphology and constantly monitor their microenvironment through the retraction and extension of fine processes (Nimmerjahn, Kirchhoff, & Helmchen, 2005). A broad range of cell surface receptors are involved in this immune surveillance function. For example, C-X3-C motif chemokine receptor 1 (CX3CR1) on microglia allows cross-talk with neurons that express C-X3-C motif chemokine ligand 1 (CX3CL1) (Harrison et al., 1998). The purinergic receptor P2RY12 aids in the detection and migration of microglia to damaged sites as first responders to injury (Haynes et al., 2006). In diseased contexts, upregulation of human leukocyte antigen (HLA) complex and inflammatory mediator expression suggests they orchestrate the dynamic process of neuroinflammation (Butovsky & Weiner, 2018).

The establishment of a unique homeostatic microglial signature is achieved through a stepwise and tissue factor-dependent differentiation process (Butovsky & Weiner, 2018). Microglia precursors arise from yolk sac hematopoiesis and colonize the brain during embryonic development, a process that is highly dependent on colony stimulating factor 1 receptor (CSF1R) (Ginhoux et al., 2010; Oosterhof et al., 2019; Rojo et al., 2019). Ligands of this receptor include the growth factors colony stimulating factor 1 (CSF1) and interleukin-34 (IL-34), which are differentially produced in the brain: CSF1 is primarily expressed by glial cells including microglia, whereas IL-34 is mainly expressed by neurons (Kana et al., 2019). The latter is an example of tissue-specific cues driving microglia differentiation, as *Il-34*^*LacZ/LacZ*^ mice show selective absence of microglia and Langerhans cells, but not of other myeloid populations (Wang et al., 2012). Another signaling pathway important for microglia identity acquisition is that of transforming growth factor-beta (TGF-β) (Butovsky et al., 2014; Gosselin et al., 2017), without which microglia fail to develop properly in mouse brain (Butovsky et al., 2014). The importance of tissue-specific cues in microglia development has been further supported by studies that showed partial acquisition of microglia characteristics by bone marrow-derived monocytes upon transplantation into brain (Bennett et al., 2018; Shemer et al., 2018).

The *in situ* identity of microglia is rapidly lost when transferred to a cell culture environment, with downregulation of homeostatic markers such as *CX3CR1* and *P2RY12* (Gosselin et al., 2017). Loss of exposure to brain-derived cues upon isolation likely contributes to this phenomenon, since the transcriptomic alteration can be partly prevented by supplementing the media with CSF1/IL-34 and TGF-β1 (Butovsky et al., 2014; Gosselin et al., 2017). From these observations derive successful generation of microglia-like cells from induced pluripotent stem cells (iPSCs) using CSF1, IL-34 and TGF-β1 (Abud et al., 2017; McQuade et al., 2018). Co-culture of primary microglia or iPSC-derived microglia (iMGL) with other brain cells has been proven to be beneficial in further reproducing *in situ* microglia characteristics (Abud et al., 2017; Grubman et al., 2020; Timmerman et al., 2022).

An additional variable in the current methods for the culture of primary microglia is the widespread use of animal-derived serum. Since its introduction in the 1950’s (Puck, Cieciura, & Robinson, 1958), fetal bovine serum (FBS) has been used to promote cell survival and growth *in vitro*, but contains bioactive molecules that might not be present in the brain (Subbiahanadar Chelladurai et al., 2021). A serum-free alternative medium for microglia has been proposed by Bohlen *et al*., who discovered that the combination of CSF1R ligand, TGF-β2 and cholesterol can sustain rodent microglia survival in serum-free conditions and promote a ramified morphology (Bohlen et al., 2017). The impact of such serum-replacement strategy on primary human microglia (hMGL) had yet to be studied in a systematic manner. The present study aims to clarify the effects of serum and growth factors supplementation of culture media on hMGL survival, immune functions and signature gene expression.

## 2. Materials and Methods

### 2.1. Cell culture reagents

A list of cell culture reagents used in this study is provided as supporting information (Supplementary table S1).

### 2.2. Microglia isolation and culture

Isolation of hMGL was carried out as previously described (Durafourt, Moore, Blain, & Antel, 2013) with slight modifications. Briefly, human brain tissues were obtained from non-malignant cases of temporal lobe epilepsy, at sites distant from suspected primary epileptic foci. Samples were digested by trypsin (Thermo Fisher Scientific) and DNase (Roche) and passed through a nylon mesh filter. Tissue homogenate was then subjected to Percoll (Sigma-Aldrich) gradient centrifugation. Further purification of microglia was performed through magnetic-activated bead sorting of CD11b^+^ cells (Miltenyi Biotec). This isolation method results in a culture of ∼97% PU.1^+^, ∼1% O4^+^ and 0% glial fibrillary acidic protein-positive (GFAP^+^) cells (data not shown). Cells were maintained at 37 °C under a 5% CO_2_ atmosphere. Unless otherwise specified, cells were cultured for six days prior to downstream experiments to allow cell adhesion to cultureware and stabilization of morphologies, a duration similar to those of other studies (Bohlen et al., 2017; Butovsky et al., 2014; Gosselin et al., 2017; Timmerman et al., 2022). Use of human cells was approved by the McGill University Health Centre Research Ethics Board. Microglia from at least one female donor and one male donor ranging between 2 and 68 years old were included in each experiment.

### 2.3. Cell viability

#### Primary microglia

Cells were stained in phosphate buffered saline (PBS) containing 1 μg/mL propidium iodide (PI) and 5 μg/mL Hoechst 33342 (Thermo Fisher Scientific). The average number of PI-live cells per condition was determined using a CellInsight CX7 High Content Screening Platform (Thermo Fisher Scientific). All conditions were assessed in triplicate.

#### iMGL

Due to the loose adherence of iMGLs to cell culture plates (Abud et al., 2017; McQuade et al., 2018), the quantification of PI-live cells was carried out using flow cytometry (Attune™ Nxt Flow Cytometer; Thermo Fisher Scientific). Doublets and debris were excluded from the analysis using appropriate forward/side scatter profiles.

### 2.4. Immunocytochemistry

Cells were fixed in 4% formaldehyde and permeabilized/blocked using PBS with 3% goat serum and 0.2% triton X-100. Cells were incubated at 4°C overnight with primary antibodies (Ki67 BD Biosciences #556003 at 1:200, IBA1 #NC9288364, Fujifilm Wako Chemicals at 1:500, PU.1 #2258, Cell Signalling at 1:200, MerTK #ab52968, Abcam at 1:1000), then with secondary antibodies and 1 μg/mL Hoechst 33342. The proportion of cells with nuclear Ki67 stain was determined using a CellInsight CX7 High Content Screening Platform. All conditions were assessed in triplicate.

### 2.5. Ribonucleic acid (RNA) -sequencing

#### Sequencing

Quality control of the RNA samples, as well as the library preparation and RNA-sequencing were performed by Genome Quebec, Montreal, Canada. A NEBNext® Single Cell/Low Input RNA Library Prep Kit (New England Biolabs) was used for library generation. RNA-sequencing was performed using an Illumina NovaSeq 6000. Canadian Center for Computational Genomic’s pipeline GenPipes (Bourgey et al., 2019) was used to align the raw files and quantify the read counts. Briefly, raw fastq files were aligned to the GRCh38 genome reference using STAR aligner (Dobin et al., 2012) with default parameters and raw reads were quantified using HTseq count (Anders, Pyl, & Huber, 2015). Transcript per million (TPM) values are provided as supporting information (Supplementary table S2).

#### Differential expression gene (DEG) analysis

Read counts were used for DEG analysis with the edgeR package (Robinson, McCarthy, & Smyth, 2009). DEGs were identified using an adjusted p-value cutoff of 0.05.

#### Enrichment analysis

Gene ontology (GO) enrichment analyses were performed using the web-based tool offered by the Gene Ontology Consortium. The Enrichr web tool (Chen et al., 2013) was used for pathway enrichment analysis using Molecular Signatures Database (MSigDB) 2020 hallmark gene sets.

#### Data visualization

Heatmaps were generated using the Python data visualization library Seaborn. Scatter plots and volcano plots were generated using the Python data visualization library Matplotlib. Principal component analysis (PCA) was carried out using the Python module sklearn and visualized using Matplotlib. Venn diagrams were drawn using Venny 2.1.0 (BioinfoGP). Histograms were generated using GraphPad Prism 8.0 software.

#### Conversion of mouse genes to human orthologs

Lists of mouse genes were converted to lists of human ortholog genes using the HUGO Gene Nomenclature Committee Comparison of Orthology Predictions search tool (Eyre, Wright, Lush, & Bruford, 2007).

### 2.6. Phagocytosis assay

Human myelin was prepared as previously described through repeated sucrose density centrifugation and osmotic shocks of white matter (Healy et al., 2016). Cells were challenged with 15 ng/mL of pHrodo™ Green (Thermo Fisher Scientific) -labelled myelin debris. After two hours, cells were counterstained with Hoechst 33342 (5 μg/mL) and total green fluorescence intensity per cell was quantified using a CellInsight CX7 High Content Screening Platform. All conditions were assessed in triplicate.

### 2.7. Measurement of cytokine and growth factor secretion

Concentrations of interleukin-1beta (IL-1β), interleukin-6 (IL-6), interleukin-10 (IL-10) and tumor necrosis factor alpha (TNFα) in cell supernatants were measured using the Human Inflammatory Cytokine Cytometric Bead Array Kit (BD Biosciences). A LEGENDplex™ HSC Myeloid Panel kit (Biolegend) was used for the measurement of the following growth factors in cell culture supernatants: CSF1, IL-34, colony-stimulating factor 2 (CSF2, also known as GM-CSF), interleukin-3 (IL-3), stem cell factor (SCF), C-X-C Motif Chemokine Ligand 12 (CXCL12) and FMS-like tyrosine kinase 3 ligand (FLT3L). Readings were done on an Attune™ Nxt Flow Cytometer.

### 2.8. Fetal astrocyte isolation and culture

Human astrocytes were isolated from second trimester fetal tissue (17–23 weeks of gestation) obtained from the University of Washington Birth Defects Research Laboratory and cultured as previously described (Kieran et al., 2022). Supernatants of cells maintained in normoxic condition or in 1% atmospheric oxygen in the presence of IL-1β (Thermo Fisher Scientific) for 24 hours were collected as described in (Kieran et al., 2022).

### 2.9. Generation of iMGL

Differentiation of iPSCs (DYR100; American Type Cell Collection) into iMGL was carried out following a previously established protocol (McQuade et al., 2018). Briefly, hematopoietic progenitor cells were generated from iPSCs using STEMdiff Hematopoietic kit (STEMCELL Technologies) and cultured in microglia differentiation medium (Table 1) supplemented with 100 ng/mL IL-34, 50 ng/mL TGF-β1 and 25 ng/mL CSF1 (Peprotech) for 25 days, following which 100 ng/mL cluster of differentiation 200 (CD200; Abcam) and CX3CL1 (Peprotech) were also added to the culture. Cells were maintained at 37 °C under a 5% CO_2_ atmosphere throughout the protocol. Resulting iMGL (day 28 and older) showed a ramified morphology (Supplementary figure 1A) and increased expression of microglia marker genes compared to iPSCs (Supplementary figure 1B). The cells stained positive for ionized calcium binding adaptor molecule 1 (IBA1), transcription factor PU.1 and Mer tyrosine kinase (MerTK) (Supplementary figure 1C).

**Table 1.** Composition of the Abud medium.

| **Component** | **Concentration** | **Source** |
| --- | --- | --- |
| Dulbecco's Modified Eagle Medium/Ham's F12 (DMEM/F12), phenol red-free | 1X | Thermo Fisher Scientific |
| B27 | 2X | Thermo Fisher Scientific |
| N2 | 0.5X | Thermo Fisher Scientific |
| GlutaMAX™ | 1X | Thermo Fisher Scientific |
| Non-Essential Amino Acids | 1X | Thermo Fisher Scientific |
| Insulin-Transferrin-Selenium (ITS-G) | 2X | Thermo Fisher Scientific |
| Monothioglycerol | 400 µM | Sigma Aldrich |
| Penicillin/Streptomycin (P/S) | 1% | Thermo Fisher Scientific |

### 2.10. Quantitative reverse transcription polymerase chain reaction (qRT-PCR)

RNA was extracted using a RNeasy mini kit (Qiagen) and reverse transcription was performed using Moloney murine leukemia virus reverse transcriptase (Thermo Fisher Scientific). Real-time polymerase chain reaction (PCR) was performed using TaqMan assays (Thermo Fisher Scientific) on a QuantStudio™ 5 real-time PCR system (Thermo Fisher Scientific). The 2^-ΔCt^ method was used to analyze the data using *glyceraldehyde 3-phosphate dehydrogenase* (*GAPDH*) and *tyrosine 3-monooxygenase/tryptophan 5-monooxygenase activation protein zeta* (*YWHAZ*) as controls.

### 2.11. Western blotting

Cells were lysed on ice in a buffer of 150 mM NaCl, 50 mM Tris-HCl pH 7.4, 1% Nonidet P-40, 0.1% sodium dodecyl sulfate (SDS) and 5 mM ethylenediaminetetraacetic acid with protease and phosphatase inhibitors (Pierce). Cell lysates were centrifuged at 500 g for 30 minutes at 4°C to remove cellular debris. Proteins (30 μg/lane) were separated on an SDS-polyacrylamide gel and transferred to a polyvinylidene difluoride (PVDF) membrane (Bio-Rad Laboratories). The membrane was immunoblotted for CSF1R (#MAB3291, R&D Systems at 1:250) and GAPDH (#G8795, Sigma Aldrich at 1:500) overnight at 4°C and then with horse radish peroxidase-linked goat anti-mouse antibodies (1:10000; Jackson Laboratory) for one hour. Bands were detected by enhanced chemiluminescence (ECL) with Clarity Max ECL substrates (Bio-Rad Laboratories) using a ChemiDoc Imaging System (Bio-Rad Laboratories). Image analysis was performed using ImageLab 6.0.1 software (Bio-Rad Laboratories).

### 2.12. Statistical analyses

Statistical analyses were performed using GraphPad Prism 8.0 software. A t-test was used to compare the mean of two groups of data. A one-way analysis of variance (ANOVA) was used to compare the mean of three or more groups of data. When the assumptions of a t-test (homoscedasticity and normal distribution) were not met, a Mann Whitney test was employed instead. When the assumptions of a one-way ANOVA (homoscedasticity and normal distribution) were not met, a Kruskal-Wallis test was used instead. P-values were adjusted using Sidak’s post hoc test and Dunn’s post hoc test following a one-way ANOVA and a Kruskal-Wallis test, respectively. A mixed-effects analysis followed by Tukey’s post hoc test was used for the comparison of multiple groups of data that depend on several independent variables. Mean and standard error of the mean (SEM) of biological replicates are plotted in all graphs unless otherwise indicated. A p-value < 0.05 was considered statistically significant.

## 3. Results

### 3.1. Brain-derived growth factors do not improve hMGL survival in serum-free culture

Traditionally, hMGL have been cultured in Minimum Eagle’s Medium supplemented with 1% glutaMAX™, 0.1% glucose and 1% penicillin/streptomycin (henceforth abbreviated MEM for simplicity) with 5% FBS to ensure cell survival (Abud et al., 2017; Durafourt et al., 2013; Healy et al., 2016). Based on findings related to the culture of rodent microglia (Bohlen et al., 2017), we first sought to replace FBS using a combination of TGF-β1, IL-34, CSF1, CD200, CX3CL1 and cholesterol, which we abbreviate to TIC_4_. In addition to the growth factors and cholesterol, both CD200 and CX3CL1 were added as brain-derived cues that are known to modulate microglia function (Abud et al., 2017; Harrison et al., 1998; Hoek et al., 2000). When hMGL survival was assessed on the sixth day of culture using PI staining, a significant reduction in survival was observed in MEM compared to MEM + 5% FBS (Figure 1A-B). Replacing FBS by TIC_4_ only partly rescued cell viability (Figure 1A-B). We hypothesized that the serum-free media used for the generation and maintenance of iMGL might be able to support the survival of hMGL. As expected, hMGL cultured in the medium formulated for iMGL generation by Abud *et al*. (Abud et al., 2017) had similar viability to those cultured in MEM + 5% FBS (Figure 1A-B). The addition of FBS, but not TIC_4_, to the Abud medium further increased cell viability (Figure 1A-B). In general, hMGL appeared more ramified in serum-free culture (Supplementary Figure S2).

**Figure 1.**
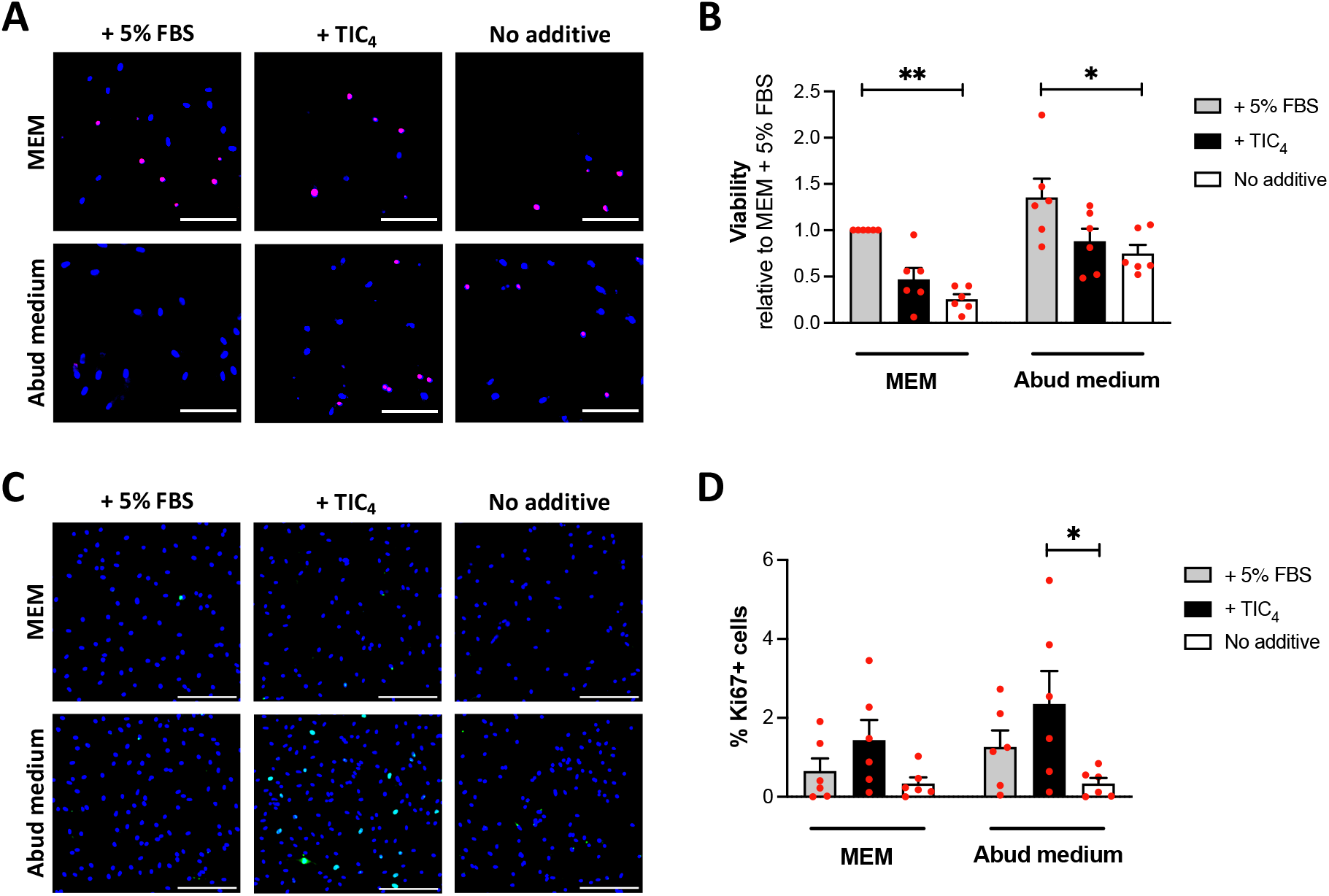
Survival and proliferation of hMGL in serum-free culture. (A) Fluorescent images of hMGL cultured for six days in various culture media stained with PI (red) and Hoechst 33342 (blue). Scale bar = 250 μm. (B) Viability of hMGL after six days in culture. A one-way ANOVA with Sidak’s post hoc test was performed. Mean +/-SEM of n=6, *p<0.05. (C) Fluorescence images of hMGL cultured for five days in various culture media and stained for Ki67 (green) and with Hoechst 33342 (blue). Scale bar = 500 μm. (D) Percentage of Ki67+ hMGL on the fifth day of culture. A one-way ANOVA with Sidak’s post hoc test was performed. Mean +/-SEM of n=6, *p<0.05.

To verify whether proliferation of hMGL contributed to the differential effect of culture media on viable cell yield assessed on day 6, Ki67 immunostaining was performed on day 5. The proportion of cells with nuclear accumulation of Ki67 was no different between MEM + 5% FBS and the serum-free Abud medium (Figure 1C-D). TIC_4_ significantly increased the proportion of nuclear Ki67+ cells only when added to the Abud medium, from ∼0% to ∼2% (Figure 1C-D). FBS supplementation did not affect cell proliferation (Figure 1C-D). Taken together, the data indicate that i) while hMGL survive in serum-free condition, FBS further enhances survival, and ii) TIC_4_ cannot replace FBS for this purpose.

### 3.2. Culture of hMGL induces a significant transcriptomic change regardless of media formulation

To determine in which culture media hMGL are most representative of their *in situ* profile, the transcriptome of *in vitro* hMGL after six days in culture was compared to that of immediately *ex vivo* hMGL from the same donors using RNAseq. Because hMGL in MEM and MEM + TIC_4_ showed poor survival, these conditions were excluded from further experiments. PCA revealed a large separation between *ex vivo* hMGL samples and all *in vitro* hMGL samples along PC1 with an overall variation of 41.6% (Figure 2A). Only small variations were observed between *in vitro* hMGL samples from different donors and from different cell culture conditions, which spread on PC2 with an overall variation of 14.5% (Figure 2A). This is indicative of a major change in cell transcriptome caused by the *ex vivo* to *in vitro* transition, which was not prevented by any of the media formulations tested. Accordingly, DEG analysis revealed that large number of genes are commonly up/downregulated by the different culture conditions compared to the *ex vivo* state (Figure 2B). Enrichment analysis carried out on the 1046 genes that were commonly upregulated in all *in vitro* conditions revealed alterations in metabolic processes such as “oxidative phosphorylation”, “fatty acid metabolism”, “mammalian target of rapamycin complex 1 (mTORC1) signaling” and “glycolysis” (Figure 2C). The 1515 genes commonly downregulated in all *in vitro* conditions were enriched in genes related to “TNF-α signaling via nuclear factor-kappa B (NF-κB)” (Figure 2C). Among them were immediate early genes previously discovered to be upregulated by the tissue dissociation procedure (van den Brink et al., 2017) including genes encoding for heat shock proteins and members of the Fos gene family (Supplementary Table S3). Removal of such genes from the analysis did not affect the outcome of the enrichment analysis (data not shown).

**Figure 2.**
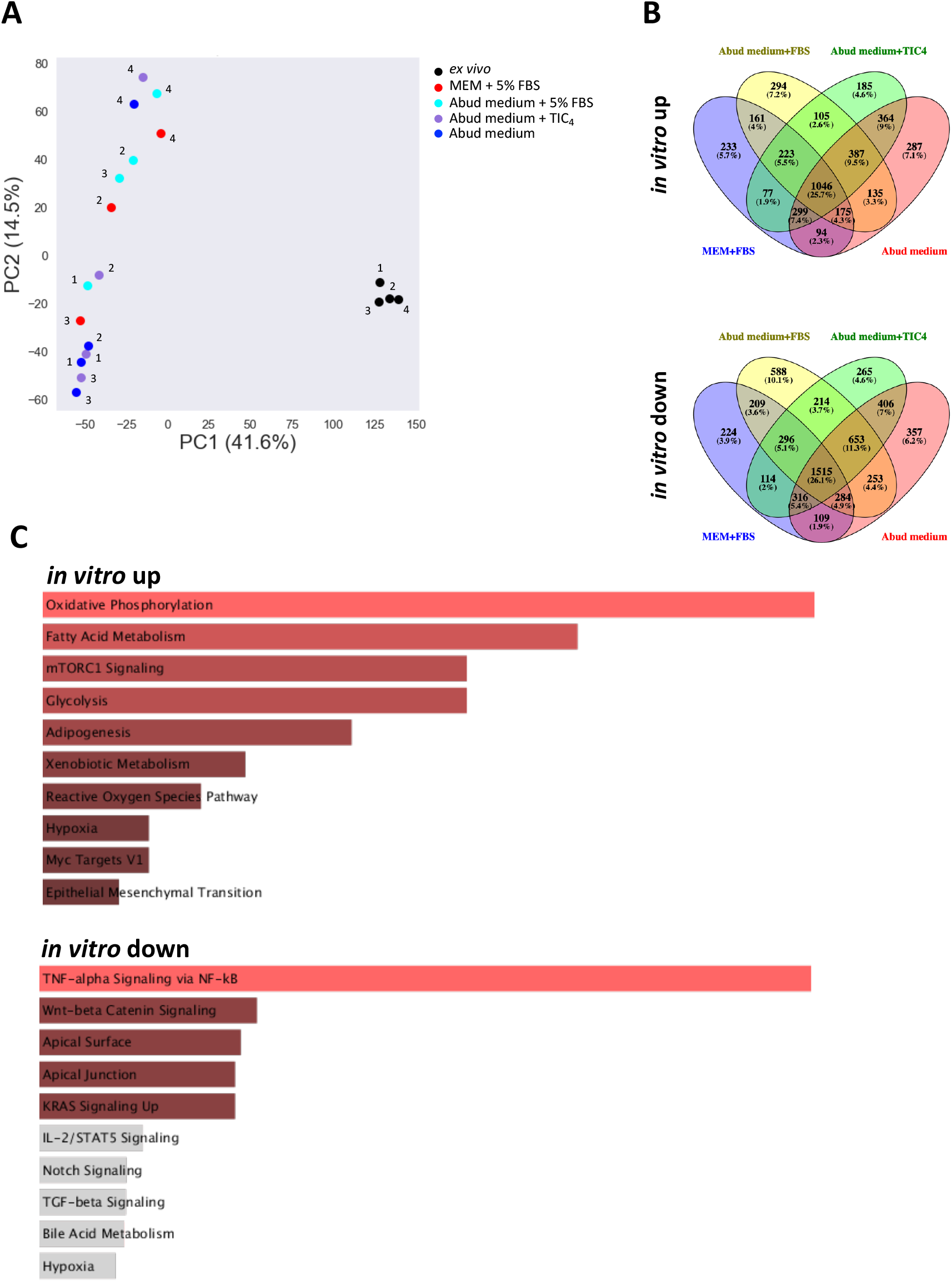
Transcriptomic alterations induced by *in vitro* maintenance of hMGL. hMGL were cultured for six days in different media or not (*ex vivo*) following isolation from brain tissues. hMGL from four donors were used. (A) PCA plot of RNA-seq data. Number denotes individual donor (1: 4-year-old male, 2: 17-year-old male, 3: 31-year-old female, 4: 15-year-old female). (B) Venn diagrams showing the number of significantly up (left diagram) or down (right diagram) -regulated genes when *in vitro* hMGL were compared to *ex vivo* hMGL. A p<0.05 was considered statistically significant in DEG analysis. (C) Results of enrichment analysis comparing the lists of genes significantly up (left table) or down (right table) -regulated *in vitro* versus *ex vivo* to the MSigDB Hallmark 2020 gene sets. A p<0.05 was considered statistically significant in DEG analysis.

In 2017, Gosselin *et al*. were the first to study the transcriptomic alteration induced in hMGL by *in vitro* maintenance (Gosselin et al., 2017). The vast majority of genes identified to be dysregulated *in vitro* (in DMEM/F12 with GlutaMAX™, FBS, penicillin, streptomycin, amphotericin B and IL-34) compared to *ex vivo* state in their study followed the same patterns of alteration in our culture conditions (Supplementary Figure S4), further supporting that media formulation has a minor impact on culture-induced changes in hMGL transcriptome. Upregulated in both studies were genes encoding chemokines, as well as stress response-related genes: sequestosome 1 (*SQSTM1*, encoding p62), peroxiredoxin 1/5 (*PRDX1/5*) and hypoxia-inducible factor 1 alpha (*HIF1A*) (Supplementary Figure S3). Downregulated in both studies were genes of the human leukocyte antigen (HLA) system and genes encoding toll-like receptors (Supplementary Figure S3).

### 3.3. FBS helps maintain microglia signature genes

As the overall impact of culture media on hMGL transcriptome was small (compared to the *ex vivo* to *in vitro* transition effect), the expression of select microglia signature genes was next evaluated. In 2014, Butovsky *et al*. identified six genes that distinguishes microglia from other myeloid and brain cells: growth arrest specific 6 (*GAS6*), protein S (*PROS1*), complement C1q A chain (*C1QA*), *MERTK, P2RY12* and G-protein coupled receptor 34 (*GPR34*) (Butovsky et al., 2014). RNAseq analysis of their expression revealed significant downregulation of most genes in Abud medium compared to *ex vivo* hMGL (Figure 3A). Surprisingly, there was no significant downregulation of those genes when cells were cultured in the presence of FBS (Figure 3A). The expression of other genes that characterize homeostatic microglia such as *CSF1R* and sialic acid binding Ig like lectin 10 (*SIGLEC10*) were also higher in Abud medium + 5% FBS compared to Abud medium alone (Figure 3B). Supplementation of the medium with TIC_4_ did not result in the same beneficial effect (Figure 3A-B). In fact, DEG analysis revealed that addition of TIC_4_ to Abud medium does not induce any significant changes in hMGL transcriptome, whereas addition of FBS resulted in 1.7% and 1.9% of genes being up and downregulated, respectively (Figure 3C). Neither FBS nor TIC_4_ supplementation had an effect on *CX3CR1*, transmembrane protein 119 (*TMEM119*), olfactomedin like 3 (*OLFML3*) and spalt like transcription factor 1 (*SALL1*) expression (Figure 3B).

**Figure 3.**
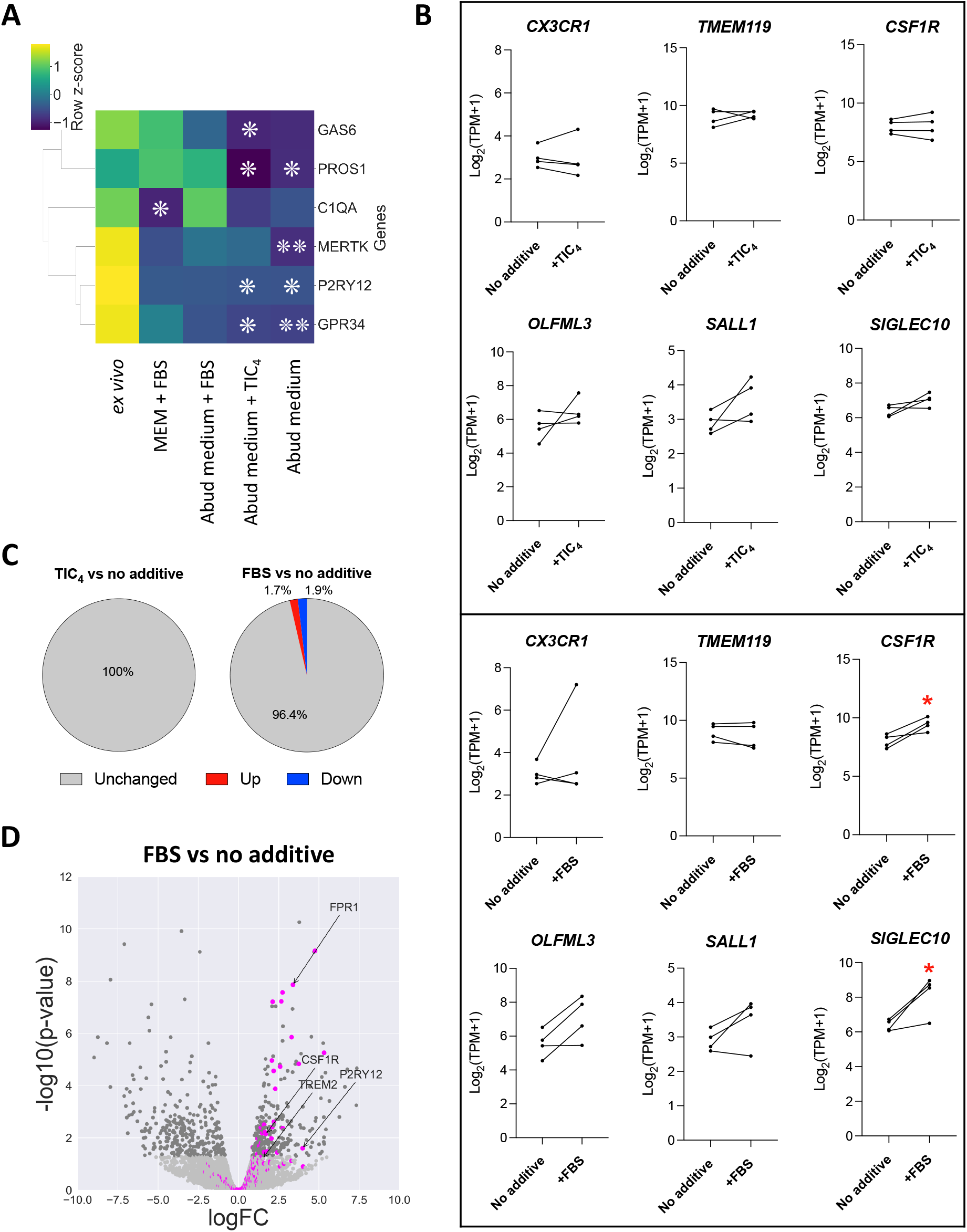
Differential effect of media formulation on microglia key signature genes expression. hMGL were cultured for six days in different media or not (*ex vivo*) following isolation from brain tissues. hMGL from four donors were used. (A) Heatmap showing the expression of microglia signature genes identified by Butovsky *et al*. (Butovsky et al., 2014). Kruskal-Wallis tests were performed, followed by Dunn’s post hoc test. *p<0.05, **p<0.01 compared to *ex vivo* hMGL. (B) Expression of select microglia markers in hMGL cultured in Abud medium supplemented with TIC_4_ or 5% FBS compared to Abud medium. T-tests were performed. *p<0.05. (C) Pie charts presenting the proportion of DEGs in Abud medium supplemented with TIC_4_ or 5% FBS, compared to Abud medium. Results are presented as percentages of the total number of genes. (D) Volcano plot of the transcriptomic change induced by 5% FBS supplementation of Abud medium. Light gray and dark gray dots depict genes for which p>0.05 and p<0.05 by DEG analysis, respectively. Fuchsia dots depict microglia signature genes that were identified by Patir *et al*. (Patir et al., 2019).

A more extensive list of genes that distinguishes microglia from other brain cells was generated by Patir *et al*. through comparison of several published transcriptomic datasets (Patir, Shih, McColl, & Freeman, 2019). DEG analysis confirmed that FBS has an effect in maintaining the expression of many of those microglia core signature genes, as they were observed to be more highly expressed in Abud medium + 5% FBS than in Abud medium alone (Figure 3D).

With the advent of single-cell transcriptomic technologies, several disease-associated microglia clusters have been identified in mouse models of disease. These include inflammatory microglia identified in LPS-injected mice (Sousa et al., 2018), “disease-associated microglia” identified in a mouse model of Alzheimer’s disease (Keren-Shaul et al., 2017) and “injury-responsive microglia” identified in lysophosphatidylcholine-induced demyelination mouse model (Hammond et al., 2019). No evidence was found to support that FBS promotes the adoption of those disease model-associated signatures (Supplementary Figure S4).

### 3.4. FBS alters the phagocytic capacity and immune responsiveness of hMGL

Because culture media had differential effect on the expression of *MERTK, PROS1* and *GAS6* with known importance in microglia phagocytic processes (Healy et al., 2016), the impact of media formulation on myelin phagocytosis by hMGL was next investigated. It was observed that hMGL have greater phagocytic capacity in Abud medium + 5% FBS, compared to any other media formulations (Figure 4A-B). Relative to Abud medium, the addition of FBS increased myelin uptake ∼7.5-fold (Figure 4A-B). This effect of FBS on enhancing myelin internalization was maintained even when FBS was removed immediately prior to the phagocytosis assay (Figure 4C), indicating that this effect is not due to the presence of opsonizing molecules in FBS. When the expression of genes related to phagocytosis was evaluated, FBS was observed to significantly increase the expression of genes encoding MerTK (Healy et al., 2016) and CD36 (Grajchen et al., 2020) with known roles in myelin phagocytosis (Figure 4D).

**Figure 4.**
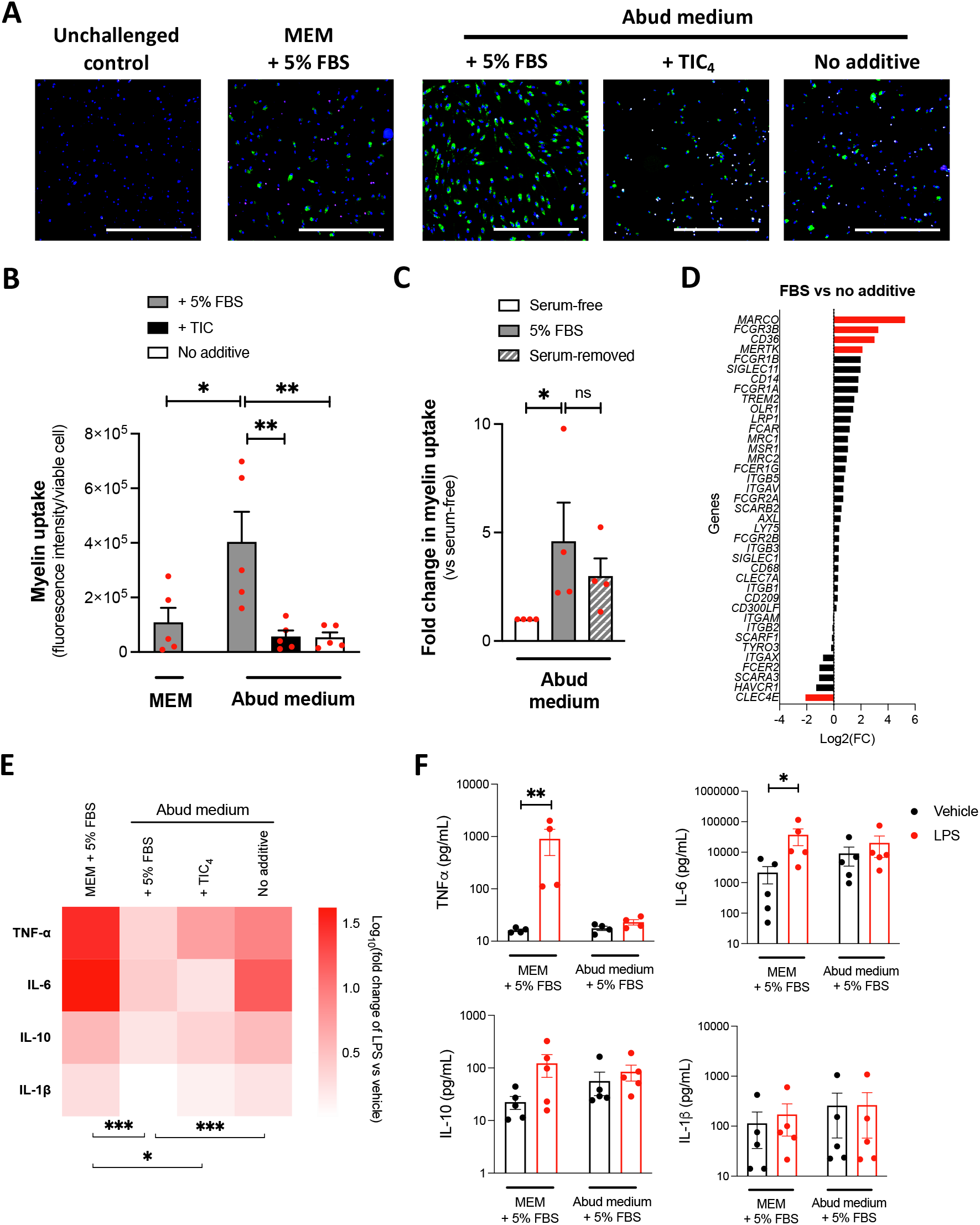
Impact of media formulation on the phagocytic and inflammatory capacity of hMGL. hMGL were cultured for six days in various culture media and (A-D) exposed or not for two hours to pHrodo Green™-labelled myelin debris or (E-F) exposed for 24 hours to vehicle or 100 ng/mL LPS. (A) Fluorescent images of internalized myelin (green) in cells stained with PI (red) and Hoechst 33342 (blue). Scale bar = 1 mm. (B) Quantification of myelin uptake normalized to the number of viable cells. A one-way ANOVA with Sidak’s post hoc test was performed. Mean +/-SEM of n=5, *p<0.05, **p<0.01. (C) Quantification of myelin uptake in cells exposed or not to FBS during myelin challenge. A Kruskal-Wallis test with Dunn’s post hoc test was performed. Mean +/-SEM of n=4, ns = non-significant, *p<0.05. (D) Transcriptional expression of membrane proteins involved in phagocytosis in hMGL cultured in Abud medium + 5% FBS compared to Abud medium (mean of n=4). Red bar denotes genes for which p<0.05 in DEG analysis. (E) Heatmap presenting the LPS-induced secretion of cytokines relative to vehicle treatment. A mixed-effects analysis followed by Tukey’s post hoc test was used. Mean of n=5, *p<0.05, **p<0.01, ***p<0.001. (F) Quantification of cytokine concentrations in cell supernatants. A Kruskal-Wallis test followed by Dunn’s post hoc test was performed. Mean +/-SEM of n=5, *p<0.05, **p<0.01.

Another important function of microglia is to elicit an inflammatory response to pathogen and danger-associated molecular patterns. The secretion of TNF-α, IL-6, IL-10 and IL-1β was measured following cell treatment with the widely used inflammatory stimulus LPS. Secretion of these cytokines was observed to be dampened in Abud medium with or without TIC_4_/FBS compared to MEM + 5% FBS, with the lowest secretory activities observed in Abud medium + 5% FBS (Figure 4E-F). Accordingly, GO term analysis suggested a difference in the expression of genes involved in immune system processes and response to stimuli in hMGL cultured in MEM + 5% FBS compared to all other media (Supplementary Table S4). Altogether, these data suggest that media formulation can greatly affect *in vitro* observations of microglia immune activities.

### 3.5. In vitro hMGL undergo a phenotypic shift characterized by resistance to CSF1R inhibition

The maintenance of microglia population in the mature brain depends on CSF1R signaling, as both pharmacological inhibition of CSF1R (Elmore et al., 2014; Zhan et al., 2020) or sequestration of CSF1/IL-34 by antibodies (Easley-Neal, Foreman, Sharma, Zarrin, & Weimer, 2019) results in microglia depletion from mouse brain. Yet, we did not observe any improvement in hMGL survivability or transcriptomic/functional changes upon supplementation of culture media with TIC_4_. This suggested that hMGL might produce and secrete CSF1 themselves, negating the need for further supplementation with CSF1R ligands. RNAseq revealed an increase in *CSF1* and a decrease in *CSF1R* expression over *ex vivo* to *in vitro* transition of the cells. The expression of IL-34 was low (log_2_(TPM+1) < 2) in both *ex vivo* and *in vitro* hMGL (Figure 5A). Accordingly, cells cultured in Abud medium secreted CSF1 (∼79 pg/mL) but not IL-34 (Figure 5B, Supplementary Figure S5A). In comparison, human astrocytes subjected to stress (hypoxia and IL-1β treatment) secreted ∼263 pg/mL CSF1 (Supplementary Figure S5B). In order to verify whether the endogenous level of CSF1 is sufficient for *in vitro* hMGL to sustain their survival in an auto/paracrine manner, cells were treated with the CSF1R inhibitor PLX3397. hMGL and iMGL both expressed CSF1R at the protein level (Figure 5C-D), however, only iMGL were susceptible to PLX3397-induced cell death upon a six-day chronic treatment (Figure 5E-F). Both hMGL and iMGL were susceptible to the toxicity of the protein synthesis inhibitor cycloheximide, which significantly decreased viability (Figure 5F). These data indicate that hMGL survival in serum-free culture is independent of CSF1R signaling.

**Figure 5.**
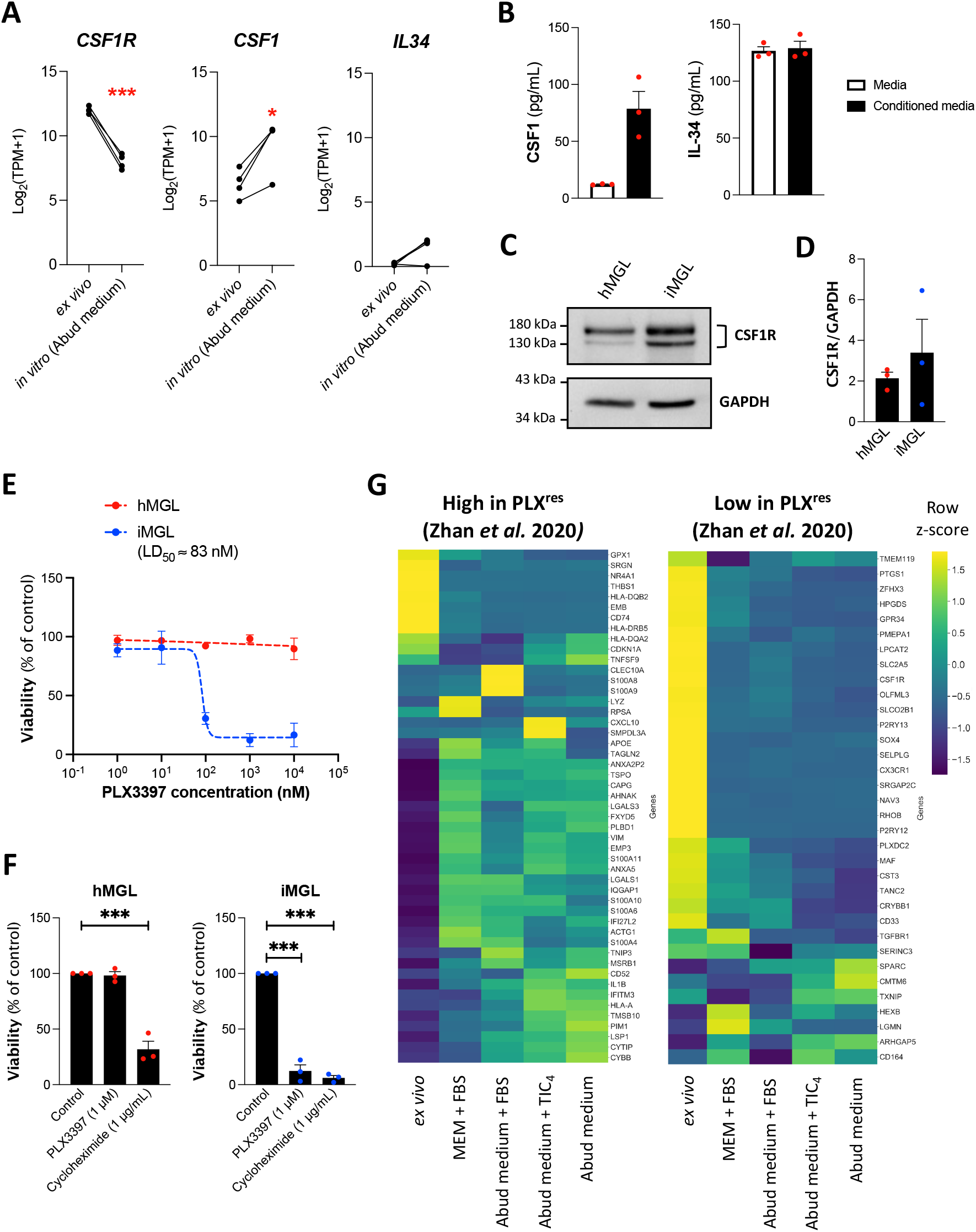
CSF1R-dependence of hMGL survival *in vitro*. hMGL were cultured for six days in different media or not (*ex vivo*) following isolation from brain tissues. (A) RNAseq assessment of *CSF1R, CSF1* and *IL-34* expression in hMGL cultured for six days in Abud medium compared to *ex vivo* hMGL obtained from the same donors. Paired t-tests were performed. n=4, *p<0.05, ***p<0.001. (B) Measurement of CSF1 and IL-34 in the supernatants of hMGL cultured for six days in Abud medium compared to unconditioned media (mean +/-SEM of n=3). (C) Immunoblotting of CSF1R and GAPDH in hMGL and iMGL cell lysates. (D) Quantification of CSF1R relative to GAPDH band intensity. Mean +/-SEM of n=3. (E) Viability of hMGL cultured in Abud medium and iMGL treated every other day with PLX3397 for 6 days. Mean +/-SEM of n=3. Dash lines represent regression curves. (F) Viability of hMGL cultured in Abud medium and iMGL treated every other day with PLX3397 or cycloheximide for six days. A one-way ANOVA followed by Dunnett’s post hoc test was performed. ***p<0.001. (G) Heatmaps showing the expression of ortholog genes identified by Zhan *et al*. (Zhan et al., 2020) to be high (left heatmap) or low (right heatmap) in PLX^res^ compared to homeostatic mouse microglia (p<0.01, |log(fold change)|>1). Genes for which log_2_(TPM+1)<2 in hMGL were excluded from the analysis.

In 2020, Zhan *et al*. identified a population of microglia in mouse brain with high expression of *Lgals3* (encoding the protein galectin-3, also called MAC2) and low expression of *Csf1r* which are resistant to the CSF1R inhibitor PLX5622 (termed PLX^res^). Gene set enrichment analysis revealed that this population differs in “TNF-α signaling via nuclear factor-kappa B”, “oxidative phosphorylation” and “mTORC1 signaling” compared to other microglia populations (Zhan et al., 2020), similarly to what is observed over the *ex vivo* to *in vitro* transition of hMGL in this study (Figure 2C). Human orthologs of most genes that are high in the murine PLX^res^ microglia were upregulated over *ex vivo* to *in vitro* transition of hMGL (Figure 5G, Supplementary figure S6). Similarly, orthologs of genes that are lowly expressed in PLX^res^ microglia were downregulated over *ex vivo* to *in vitro* transition of hMGL (Figure 5G, Supplementary figure S6). These were true regardless of cell culture media formulations (Figure 5G, Supplementary figure S6) and suggests that hMGL *in vitro* adopt a phenotype that allows for survival in the absence of CSF1R signaling, thereby reducing the need for supplementation with CSF1R ligands.

## 4. Discussion

Microglia are involved in a plethora of processes in both health and disease. The secretion of trophic factors, as well as phagocytic clearance of apoptotic cells and redundant synapses are critical for proper brain network formation (Schafer & Stevens, 2015). Immunohistochemical staining and brain imaging of patients’ brains have evidenced an inflammatory activation of microglia in several neuroinflammatory and neurodegenerative diseases (Butovsky & Weiner, 2018). In addition, genetic associations have been established between genes enriched in microglia and the risk of developing diseases such as Alzheimer’s disease (Lambert et al., 2013). It is imperative that the *in situ* microglia transcriptomic signature is faithfully replicated *in vitro* to facilitate the accurate modeling of microglia function, to expand our fundamental understanding of these cells, and to accelerate the therapeutic targeting of microglia for the treatment of neurologic diseases. In this study we investigated culture media formulation as a variable in replicating hMGL *in situ* characteristics, with a focus on how FBS and brain-derived cues could impact hMGL transcriptomics and immune activities.

By comparing the transcriptome of *in vitro* hMGL to *ex vivo* hMGL from the same donors, an unexpected beneficial effect of FBS as a culture supplement was revealed in this study. Not only did FBS supplementation improve cell survival and enhance phagocytic capacity as previously observed in rodent microglia (Bohlen et al., 2017; Fourgeaud et al., 2016), it also helped maintain expression of microglia signature genes *in vitro*. The observed increased phagocytic activity of hMGL cultured in the presence of FBS correlated with higher expression of genes encoding MerTK as well as GAS6 and PROS1, natural ligands aiding MerTK recognition of phagocytic targets (Tondo, Perani, & Comi, 2019). Supplementation of cell culture media with a mixture of cytokines containing CSF1, IL-34 and TGF-β1 induced proliferation of hMGL, but had minor impact on microglia signature genes expression, immune functions and survival over short term culture. Further characterization of hMGL in culture confirmed that these cells secrete CSF1 but do not depend on CSF1R signaling to survive. This was accompanied by a transcriptomic shift marked by decreased *CSF1R* expression compared to the *ex vivo* state.

Microglia pool in the postnatal brain is maintained through a balance between cell death and self-renewal with a slow turnover (Askew et al., 2017; Réu et al., 2017). Studies using pharmacological inhibitors (PLX3397 or PLX5622) demonstrated that this maintenance is dependent on CSF1R signaling in mice (Elmore et al., 2014; Zhan et al., 2020). Here it was observed that hMGL in culture are resistant to pharmacological inhibition of CSF1R. Cells showed decreased expression of *CSF1R* compared to the *ex vivo* state and shared transcriptomic similarities with a PLX5622-resistant microglia population identified in the mouse brain (Zhan et al., 2020). It is unlikely that selective survival of a pre-existing CSF1R-independent population of hMGL is expanding in culture, as addition of TIC_4_ did not increase cell yield nor did it prevent a decrease in CSF1R expression. Rather, it is likely that hMGL in culture adopt a cellular phenotype characterized by a reduced reliance on signaling through the CSF1R. This has profound implications in the study of CSF1R as a therapeutic target. Adult-onset leukoencephalopathy with axonal spheroids and pigmented glia (ALSP) is a fatal disease caused by inactivating mutations of CSF1R and is characterized by low number, dysmorphic microglia in the brain. Aside from its pro-survival and differentiation effect, the mitogenic effect of CSF1R signaling has been linked to exacerbated inflammation in neurological diseases such as Alzheimer’s disease (Gomez-Nicola & Perry, 2016; Hu et al., 2021) and multiple sclerosis (Hagan et al., 2020). Titrated modulation of microglia density through CSF1R targeting has therefore been tested in animal models as a promising therapeutic strategy (Green, Crapser, & Hohsfield, 2020). The lack of hMGL reliance on CSF1R to survive in culture prevents the validation of microglia depletion strategy in human models. On the contrary, iMGL depended on CSF1R signaling to survive and might be a suitable model to screen for therapeutics that can modulate microglia density.

Bohlen *et al*. previously observed poor survival of rodent microglia in serum-free culture, which could be significantly improved by supplementing the cell culture medium with a combination of CSF1/IL-34, TGF-β and cholesterol (TIC) (Bohlen et al., 2017). On the contrary, hMGL could be successfully cultured in serum-free medium without the addition of TIC_4_. The discrepancy in our observations could stem from interspecies differences and/or differences in basal media used. Several crucial interspecies differences have so far been detected between rodent and primate microglia. Differences not only relate to their transcriptome, but also in the way cells react to insults, such as morphological alterations and nitric oxide synthesis in response to interferon/LPS treatment (Geirsdottir et al., 2019; Gosselin et al., 2017; Healy, Yaqubi, Ludwin, & Antel, 2019; Owen et al., 2017). Survival requirements in growth factors, hormones and/or metabolites might represent yet another distinctive difference between microglia of various species. Supporting this notion, Bohlen *et al*.’s medium + TIC fails to sustain the initial adhesion and survival of rhesus monkey microglia in culture (Timmerman et al., 2022). It should be noted that TIC_4_ did increase hMGL proliferation, an effect that was too modest to be reflected on overall cell yield over a short-term culture period. The use of TIC_4_ or some of its components might therefore be beneficial in increasing hMGL yield over long-term culture.

When the transcriptomic difference of *in vitro* versus *ex vivo* murine microglia was first described, it was shown that TGF-β1 partly prevents the culture-induced loss of microglia signature gene expression (Butovsky et al., 2014). Robust increases in *Gpr34, P2ry12, Olfml3* and *Sall1*, among other genes, was observed in TGF-β1-supplemented culture (Butovsky et al., 2014). The effect of TGF-β1 supplementation on microglia transcriptome was observed to be much more modest in studies using hMGL (Gosselin et al., 2017) and rhesus monkey microglia, with the latter even showing a TGF-β1-induced decrease in *OLFML3* expression (Timmerman et al., 2022). Similarly, TIC_4_ was not observed to have any significant effect on the transcriptome of hMGL cultured in Abud medium. FBS was surprisingly better than TIC_4_ at maintaining key gene expression in hMGL, such as that of *MERTK* and *P2RY12*. With over 1800 proteins and 4000 metabolites (Subbiahanadar Chelladurai et al., 2021), determining which component(s) of FBS is responsible for the maintenance of microglia core genes would unfortunately require tremendous time and resources. We found no advantage of using serum-free media over the conventional MEM + 5% FBS, and no media formulation tested in this study fully recapitulated the *ex vivo* transcriptome of hMGL. Specific media formulations did have a profound effect on functional readouts commonly used when assessing microglia activity, raising the question as to whether the selection of culture media should be tailored depending on the specific biological processes under investigation.

Recent efforts to more accurately model human brain tissue has led to the optimization of co-culture and higher dimensional culture systems, circumventing the problem of environment deprivation-induced changes in microglia transcriptome (Abud et al., 2017; Grubman et al., 2020; Timmerman et al., 2022). Nevertheless, microglia monoculture remains essential for the study of cell autonomous effects of genetic manipulations and the testing of potential therapeutics. This study revealed the importance of choosing appropriate culture media in evaluating key functional aspects of microglia. This work also highlights the usefulness of complementary models such as iMGL to study certain molecular pathways (e.g. CSF1R signaling pathway) that are inactive in cultured hMGL.

## Acknowledgements

We thank lab members Manon Blain, Carol Chen, Qiao-Ling Cui, Geneviève Dorval, Nicholas Kieran, Andrea Krahn, Florian Pernin, Wolgang Reintsch and Julien Sirois for experimental/technical assistance. This study was funded by the Montreal Neurological Institute New Investigator Fund. Marie-France Dorion received financial supports from the Canadian Institutes of Health Research (CIHR).

