## Supplementary information for "Systematic comparison of culture media uncovers phenotypic shift of human microglia defined by reduced reliance to CSF1R signaling"

**Supporting information**

**Supplementary table S1: List of cell culture reagents**

**Supplementary table S2: RNAseq data in TPM values**

**Supplementary table S3: List of genes up and downregulated *in vitro* compared to *ex vivo***

**Supplementary table S4: GO term analysis**

**Supplementary figure S1: Characterization of iMGL**

**Supplementary figure S2: Impact of media formulation on microglia morphology**

**Supplementary figure S3. Expression of genes identified to be up or downregulated in hMGL in culture by Gosselin *et al.***

**Supplementary figure S4. Effect of FBS supplementation on hMGL expression of "activated" microglia signature genes**

**Supplementary figure S5. Secretion of growth factors by glial cells in culture**

**Supplementary figure S6. Overlap between the transcriptomic signature of $PLX^{res}$ and the transcriptomic changes over *ex vivo* to *in vitro* transition of hMGL**

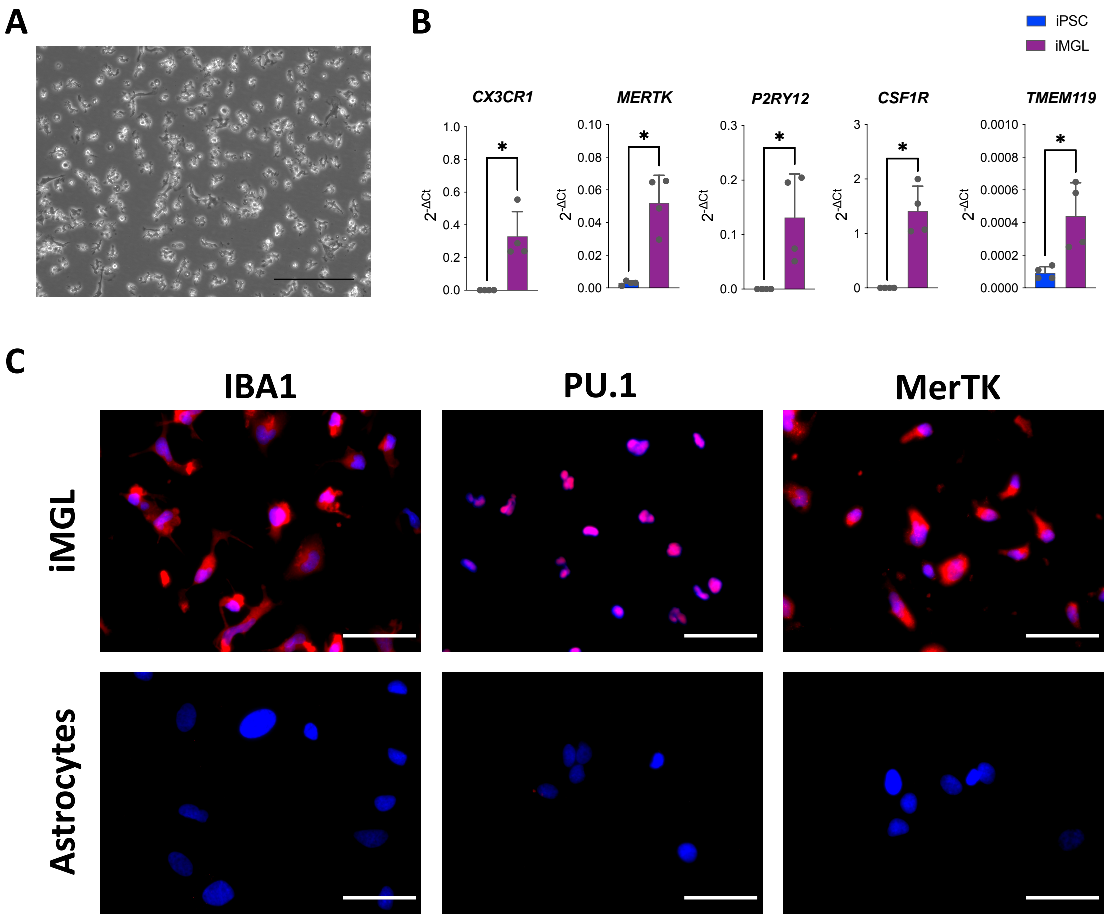


**Supplementary figure S1. Characterization of iMGL.** (A) Phase contrast image of iMGL (scale bar = 150 µm). (B) Expression of microglia marker genes in iPSC and iMGL assessed by qRT-PCR. Data are presented as $2^{-\Delta Ct}$ values obtained using *GAPDH* and *YWHAZ* as controls. A Mann-Whitney test was performed (n=4, *p<0.05). (C) Fluorescent images of iMGL and fetal astrocytes immunostained for IBA1, PU.1 or MerTK (red) and counterstained with Hoechst 33342 (blue) (scale bar = 50 µm).

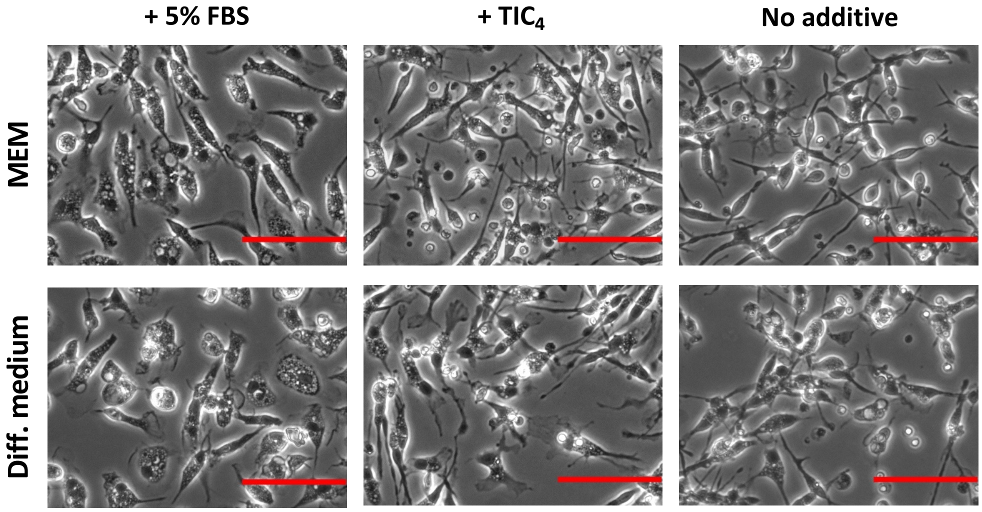


**Supplementary figure S2. Impact of media formulation on microglia morphology.** Phase contrast images of hMGL cultured in various media for six days. Scale bar = 150 μm. All images were captured at high density areas and are not representative of overall cell densities.

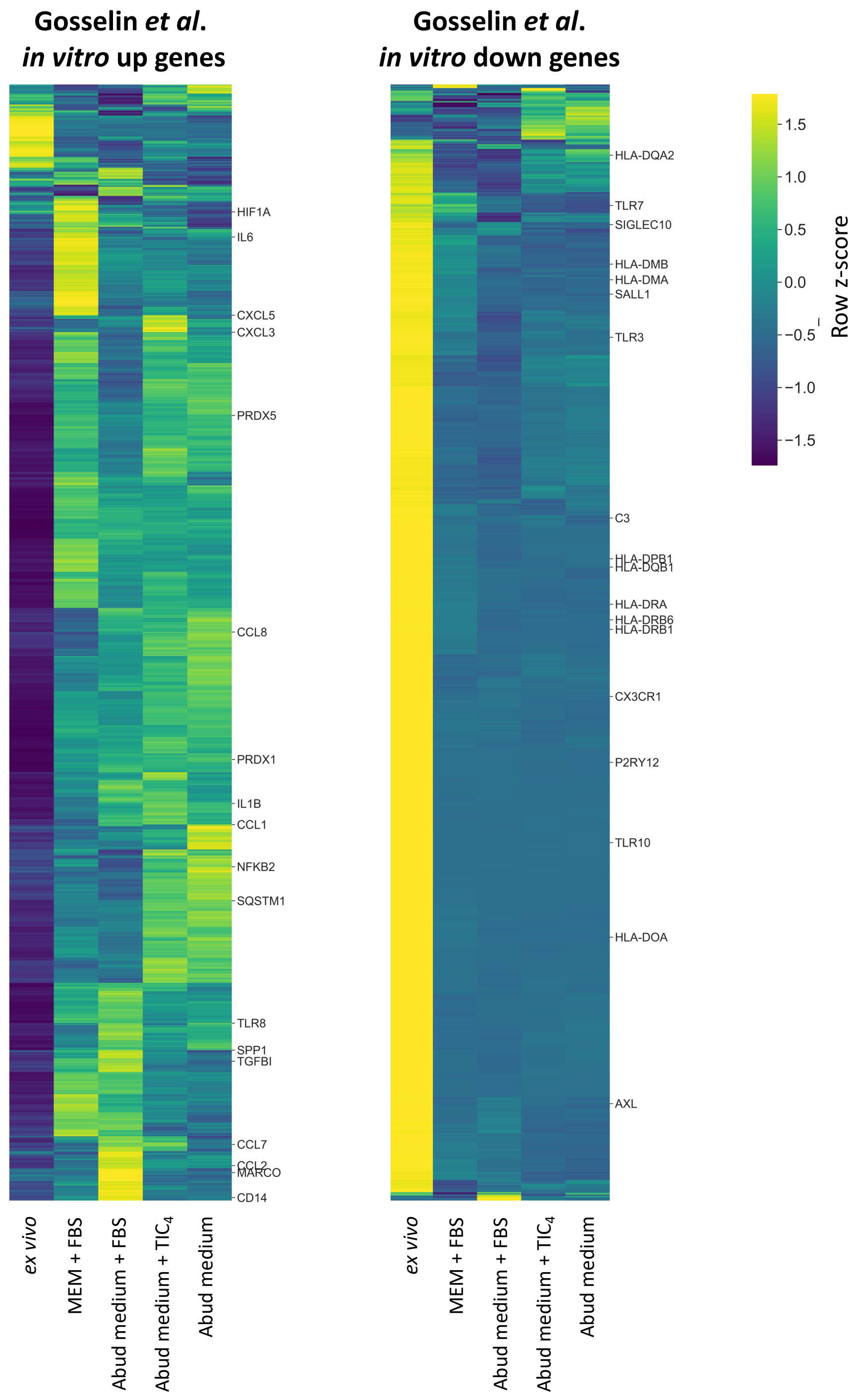


**Supplementary figure S3. Expression of genes identified to be up or downregulated in hMGL in culture by Gosselin *et al.*** hMGL were cultured for six days in different media or not (ex vivo) following isolation from brain tissues. Heatmaps show the expression of genes identified by Gosselin *et al*. to be significantly up (left heatmap) or down (right heatmap) -regulated in *in vitro* hMGL (7-day culture) compared to *ex vivo* hMGL. $p<0.01$

and $|\log(\text{fold change})|>1$ in DEG analysis was considered significant. Genes for which $\log_2(\text{TPM}+1)<2$ were excluded from the plots.

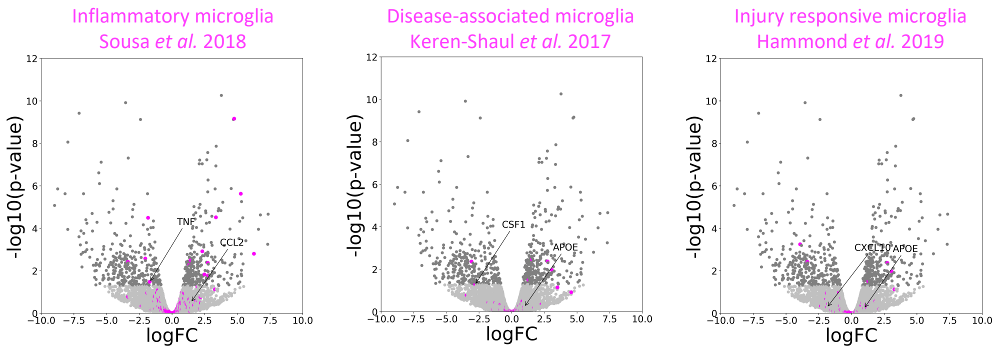


**Supplementary figure S4**. **Effect of FBS supplementation on hMGL expression of "activated" microglia signature genes.** Volcano plots of hMGL transcriptome following a six-day culture in Abud medium supplemented with 5% FBS compared to the non-supplemented counterpart. Light gray and dark gray dots depict genes for which $p>0.05$ and $p<0.05$ by DEG analysis, respectively. Fuchsia dots depict signature genes identified in microglia from LPS-injected mice (left plot, Sousa *et al.*, 2018), APP/PS1 mice (middle plot, Keren-Shaul *et al.*, 2017) and lysophosphatidylcholine-injected mice (right plot, Hammond *et al.*, 2019). FC = fold change.

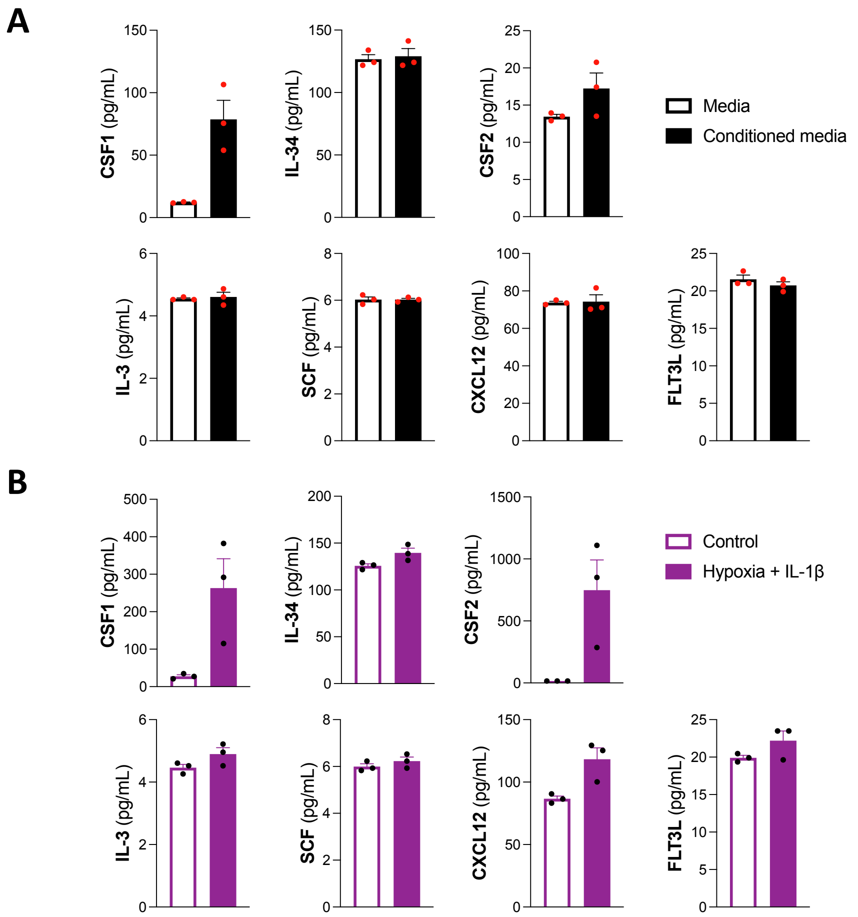


**Supplementary figure S5. Secretion of growth factors by glial cells in culture.** Growth factors measured in supernatants of (A) hMGL cultured for six days compared to cell-free wells (Mean +/- SEM of n=3), (B) fetal astrocytes subjected or not to hypoxia (1% $O_2$) and IL-1β treatment (1 ng/mL) for 24 hours (Mean +/- SEM of n=3).

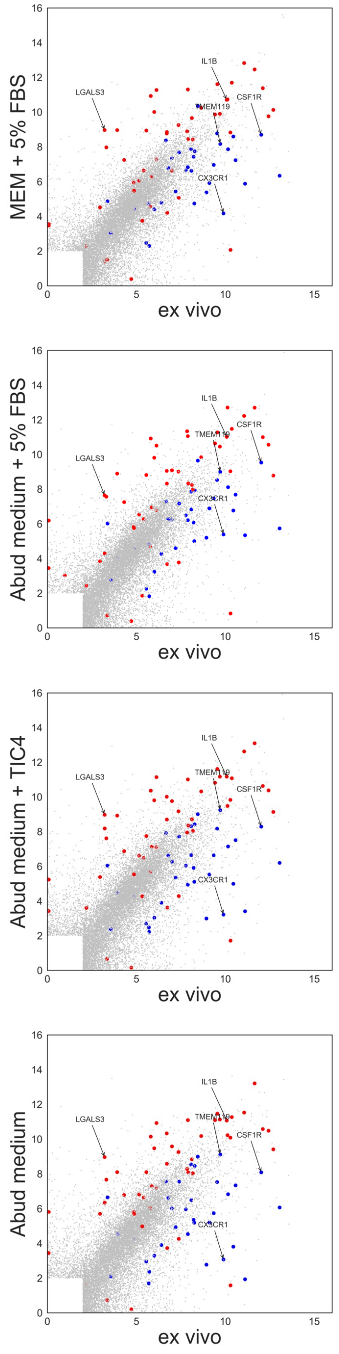


**Supplementary figure S6. Overlap between the transcriptomic signature of PLX**$^{res}$ **and the transcriptomic changes over *ex vivo* to *in vitro* transition of hMGL.** hMGL were cultured for six days in different media or not (*ex vivo*) following isolation from brain tissues. Scatter plots show the transcriptional alterations occurring over *ex vivo* to *in vitro* transition of hMGL. Ortholog genes that were observed to be high ($p<0.01$, log(fold change)>1) and low (p<0.01, log(fold change)<-1) in $PLX^{res}$ compared to homeostatic murine microglia are highlighted in red and blue, respectively. Genes for which $log_2$(TPM+1)<2 in hMGL were excluded from the plots.
